# Type I and III interferons disrupt lung epithelial repair during recovery from viral infection

**DOI:** 10.1101/2020.05.05.078360

**Authors:** J. Major, S. Crotta, M. Llorian, T. M. McCabe, H. H. Gad, R. Hartmann, A. Wack

## Abstract

Excessive cytokine signalling frequently exacerbates lung tissue damage during respiratory viral infection. Type I and III interferons (IFN-α/β and IFN-λ) are host-produced antiviral cytokines and currently considered as COVID-19 therapy. Prolonged IFN-α/β responses can lead to harmful proinflammatory effects, whereas IFN-λ mainly signals in epithelia, inducing localised antiviral immunity. Here we show that IFN signalling interferes with lung repair during influenza recovery, with IFN-λ driving these effects most potently. IFN-induced p53 directly reduces epithelial proliferation and differentiation, increasing disease severity and susceptibility to bacterial superinfections. Hence, excessive or prolonged IFN-production aggravates viral infection by impairing lung epithelial regeneration. Therefore, timing and duration are critical parameters of endogenous IFN action, and should be considered carefully for IFN therapeutic strategies against viral infections like influenza and COVID-19.

**One Sentence Summary:** A novel IFN-mediated mechanism of immunopathology during respiratory virus infection by interference with lung tissue repair.

## Main Text

During infection with respiratory viruses, disease severity is linked to the destruction of epithelia in the lung. This occurs as a result of both cytopathic viral effects in host cells, and damage generated by the ensuing inflammatory immune response. Loss of airway and lung epithelia can result in clinical complications, including lung failure, acute respiratory distress syndrome (ARDS), pneumonia, and increased susceptibility to bacterial superinfections by pathogens like *Streptococcus pneumoniae* (*S. pneumoniae*) owing to a loss of epithelial barrier function.

Restoration of damaged epithelial tissues is therefore paramount in order to maintain lung function and barrier protection. The lung epithelium is largely quiescent during steady state, yet has a remarkable capacity to replenish following damage. Several types of progenitor cells can contribute to the regeneration of the respiratory epithelium. Trp63^+^Krt5^+^ basal cells in the upper airways and Scgb1a1^+^ secretory cells in conducting airways repopulate epithelial cell populations following depletion in mice (*1, 2*). Surfactant producing type II alveolar cells (AT2) cells proliferate to reform damaged alveoli, whilst also functioning as stem cells through the differentiation into type I alveolar cells (AT1) (*3, 4*). In situations of severe damage, including influenza virus-induced injury, alveolar repair is supported by the emergence and expansion of Krt5^+^ basal cells from the airways (*5–8*).

Interferons (IFNs) represent a crucial component of the innate immune defence against viral infections. Type I and III IFNs (IFN-α/β and IFN-λ, respectively) are induced upon viral recognition, and function by triggering transcription of an overlapping set of interferon-stimulated genes (ISGs), conferring an antiviral state in infected and bystander cells (*9, 10*). Owing to widespread expression of the type I IFN receptor (IFNAR) in immune cells, IFN-α/β responses can result in immunopathology during viral infections, including influenza virus and severe acute respiratory syndrome coronavirus (SARS-CoV) (*11–14*). In contrast, the receptor for IFN-λ (IFNLR) is restricted mainly to epithelia at barrier sites like the gut and lung, with expression in the immune system largely restricted to neutrophils in mice (*15–17*). IFN-λ responses are therefore often characterised by their ability to confer localised antiviral protection at the site of infection, without driving damaging proinflammatory responses associated with IFN-α/β. Both type I and III IFNs are currently being tested in clinical trials as antiviral therapeutics against SARS-CoV-2 (*18–20*), despite some scepticism owing to IFN-α/β-associated pathogenic effects (*21*). In addition to their antiviral and proinflammatory activity, IFNs exert antiproliferative and proapoptotic functions, and have a long history in antitumour therapies (*22*). Despite a growing understanding of the causes of immunopathology during infection with respiratory viruses, little is known concerning the direct effect that IFN responses have on lung epithelial repair.

### Temporal overlap of antiviral IFN responses and lung epithelial repair following influenza virus infection

To induce acute lung damage, C57BL/6 (B6) wild type (WT) mice were infected with a sublethal dose of the H3N2 influenza virus strain X31. Influenza virus matrix gene expression peaked on day 2 post infection following viral replication, before viral clearance was reached on day 12 post infection (fig. S1A). Infection resulted in increasing morbidity and weight loss (fig. S1B), accompanied by significant immune cell infiltration and lung damage as determined by histology (fig. S1C, left, and S1D). Recovery from infection (~day 8 post infection; fig. S1B) coincided with the onset of epithelial regeneration (fig. S1C, right). To further investigate the dynamics of lung repair following influenza virus infection, epithelial cells were harvested for analysis of proliferation by flow cytometry, using the marker of recent cell proliferation Ki67 (gating strategy in fig. S2). During steady-state conditions, type II alveolar epithelial cells (AT2; EpCam^+^MHCII^+^CD49fl^ow^) (*3, 4, 23*) have a low rate of turnover, remaining largely quiescent (Fig. 1A). However, following influenza virus-induced lung damage, AT2 cells undergo rapid proliferation starting at ~day 7 post infection, correlating with mouse recovery and weight gain of mice (Fig. 1A and fig. S1B).

**Fig. 1.**
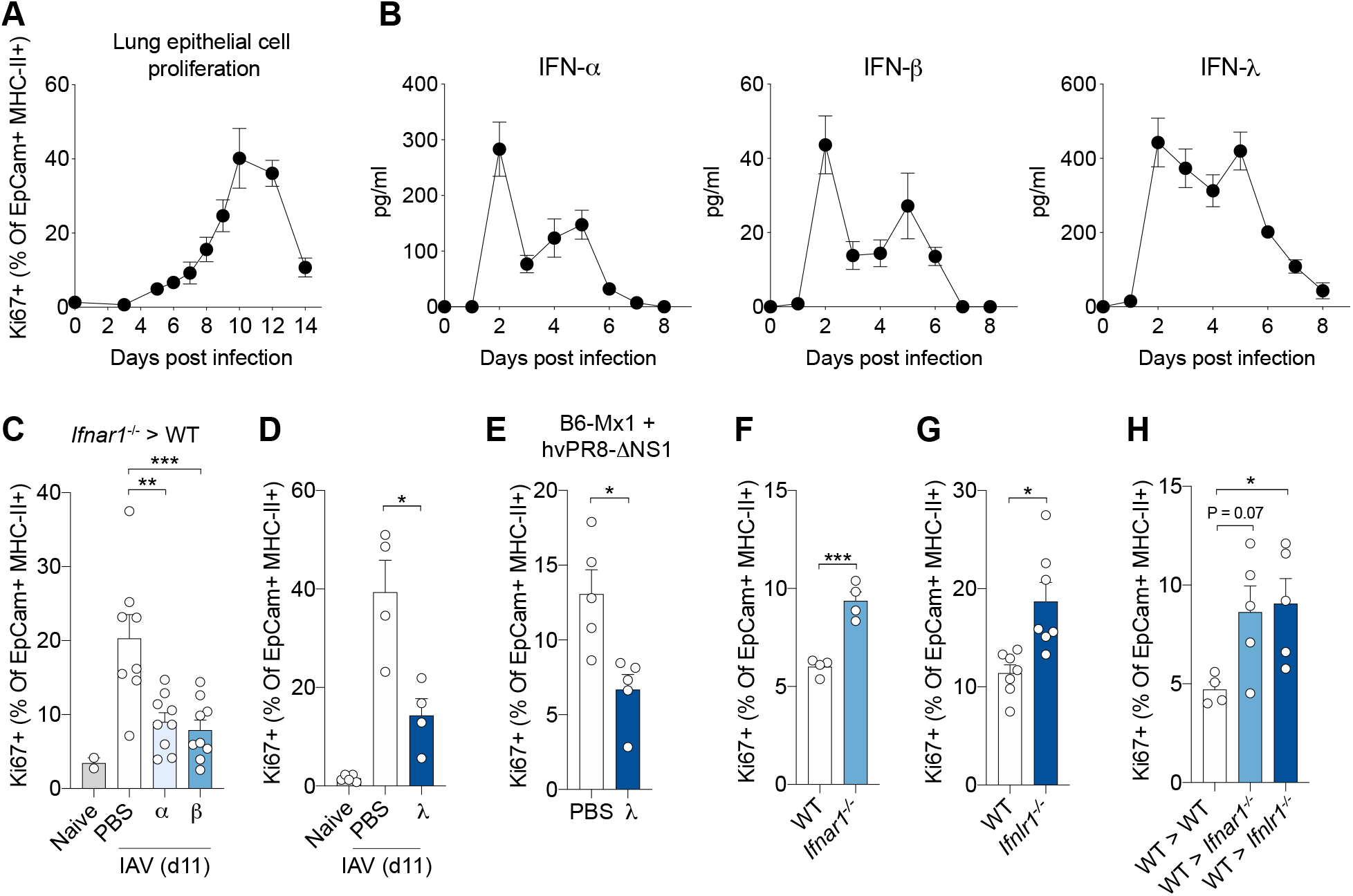
Type I and III IFNs reduce epithelial cell proliferation during lung repair. (**A** and **B**) Mice were infected with 10^4^ TCID_50_ X31 (H3N2) influenza virus in 30 μl intranasally. (A) Proliferating (Ki67^+^) AT2 cells (EpCam^+^MHCII^+^CD49f^low^) were measured by flow cytometry (n = 5), and (B) type I and III IFN levels were detected in BALF (n = 4), on indicated days post infection. (**C** and **D**) X31 infected mice were administered IFNs every 24 hours (on days 7 to 10 post infection), and proliferating (Ki67^+^) AT2 cells (EpCam^+^MHCII^+^CD49f^low^) were measured by flow cytometry on day 11 post infection. (**C**) Lethally irradiated WT mice were injected with *Ifnar1*^-/-^ bone marrow cells. Following reconstitution, influenza virus-infected chimeric mice were treated with PBS control (n = 8), IFN-α (n = 9), or IFN-β (n = 9). Naïve controls are uninfected, untreated bone marrow chimeric mice (n = 2). (**D**) Infected WT mice were treated with IFN-λ (n = 4) or PBS control (n = 4). Naïve controls are uninfected, untreated WT mice (n = 5). (**E**) B6-Mx1 mice were infected with 2.5 x 10^3^ TCID_50_ hvPR8-ΔNS1 (H1N1) and treated with IFN-λ (n = 4) or PBS control (n = 4) (IFN treatment and lung analysis was performed as for C and D). (**F** to **H**) Lungs from X31 infected WT (n = 4 to 7), (**F**) *Ifnar1*^-/-^ (n = 4), (**G**) *Ifnlr1*^-/-^ (n = 7), and (**H**) BM chimeric mice (n = 4 to 5) mice were harvested and proliferating (Ki67^+^) AT2 cells were measured by flow cytometry on day 8 post infection. All data are representative of at least two independent experiments. Data are shown as mean ± s.e.m. and statistical significance wasassessed by one-way ANOVA with Dunnett’s post-test (C, D and H), or unpaired two-tailed Student’s t-test (E to G). ns, not significant (P >0.05); \**P* ≤ 0.05, \*\**P* ≤ 0.01, \*\*\**P* ≤ 0.001.

To compare the dynamics of epithelial recovery with production of IFN in our infection model, we collected bronchoalveolar lavage fluid (BALF) at different timepoints throughout influenza virus infection (Fig. 1B). IFN subtypes (IFN-α, -β and -λ) were produced rapidly, peaking on day 2 post infection (Fig. 1B). The magnitude of IFN-λ production was significantly greater than that of IFN-α/β, both in concentration and duration. Whilst IFN-α/β production declined following the day 2 peak, high levels of IFN-λ production were maintained. Importantly, only IFN-λ was detected on day 7/8 post infection, coinciding with the onset of epithelial recovery (Fig. 1A and B). Together, these data show that following influenza virus infection, IFN signalling (in particular IFN-λ induced) is ongoing throughout the onset of epithelial proliferation and lung repair.

### IFN signalling during recovery from influenza virus infection impairs lung epithelial cell proliferation

Next, to understand the effect of IFNs on lung epithelial repair, mice were treated with IFN during recovery from influenza virus infection (day 7 to 10 post infection) (fig. S3A). To compare the effects of type I and III IFN subtypes at equal biological potency, IFN-α, -β and -λ subtypes were titrated on murine airway epithelial cell (AEC) cultures and analysed for the induction of antiviral ISGs (fig. S4A). Dose-response curves were then generated to obtain an EC50 for each subtype, in order to determine a conversion factor for equipotency between IFN-α, -β and -λ (fig. S4B). In order to study the effects of IFN treatment specifically on epithelial cells, we generated irradiation bone marrow (BM) chimeras where WT recipient mice were given *Ifnar1*^-/-^ bone marrow cells, thus restricting IFNAR expression to the stromal compartment.

In chimeric mice, both IFN-α and -β treatments significantly reduced the proliferation of AT2 cells on day 11 post influenza virus infection, compared to PBS treated control mice (Fig. 1C). This reduction was independent of changes in viral burden, as influenza virus matrix gene expression remained unchanged following IFN treatment (fig. S3B). We also observed a reduction in the proliferation of AT2 cells in WT mice following IFN-λ treatment (Fig. 1B), again independently from changes in viral load (fig. S3C). To exclude the possibility of IFN-λ signalling in neutrophils, we treated WT mice with a monoclonal antibody against Ly6G (clone 1A8) (fig. S3D). Even after depletion of neutrophils, we observed a significant reduction in AT2 cell proliferation following IFN-λ treatment (fig. S3E). A caveat when using inbred mouse strains in the context of influenza virus infection is the lack of a functional Mx1 protein; a crucial IFN-inducible influenza virus restriction factor in both mice and humans (*24*). We therefore infected mice that express functional Mx1 alleles (B6-Mx1) with the highly virulent influenza virus strain hvPR8-ΔNS1, for a more clinically relevant influenza model. IFN-λ treatment also significantly reduced lung epithelial proliferation in the presence of functional Mx1 (Fig. 1E).

We next sought to determine the role of endogenous IFN during lung repair, using *Ifnar1*^-/-^ and *Ifnlr1*^-/-^ mice. AT2 cells were analysed on day 8 post influenza virus infection; the time of overlap between IFN signalling and epithelial cell proliferation (Fig. 1A and B). Both *Ifnar1*^-/-^ and *Ifnlr1*^-/-^ mice had improved AT2 cell proliferation, compared to WT controls (Fig. 1F and G). This was dependent on IFN signalling specifically through the epithelium, as receptor deficiency in the stromal compartment alone was sufficient to increase lung epithelial cell proliferation (Fig. 1H). Improved proliferation was independent of major changes in viral burden, with only a slight increase in influenza virus matrix gene expression observed in *Ifnar1*^-/-^ mice on day 2 post infection, and no change in *Ifnlr1*^-/-^ mice, compared to WT controls (fig. S5A). Viral control in individual IFN receptor knockout mice is likely unaffected due to redundancy between type I and III IFN antiviral responses in epithelial cells (*25, 26*).

Despite type I and III IFN redundancy in viral control (fig. S5A), the lack of redundancy in antiproliferative IFN responses, with both *Ifnar1*^-/-^ and *Ifnlr1*^-/-^ mice displaying enhanced epithelial proliferation (Fig. 1F to H), led us to further interrogate the phenotype. As IFNAR signalling has previously been shown to be important for the production of IFN-λ during influenza virus infection (*27, 28*), we measured IFN production in the BALF of IFN receptor knockout mice. Consistent with previous findings (*28*), we observed a significant reduction in IFN-λ (and indeed IFN-α/β) production in *Ifnar1*^-/-^ mice compared to WT, yet we saw little change in IFN-α/β levels in *Ifnlr1*^-/-^ mice (fig. S5B). These findings suggest that the improved epithelial proliferation observed in *Ifnar1*^-/-^ mice, could be attributed to a reduction in IFN-λ. Reduced IFN production in *Ifnar1*^-/-^ mice is linked with steady state priming defects in these mice owing to the absence of tonic IFNAR activation in immune cells (*29*). To circumvent this, we used an αIFNAR monoclonal antibody during influenza virus infection (MAR1-5A3) administered only from the time of infection (day 0). αIFNAR treatment maintained steady state priming required for IFN-λ production (fig. S5C), yet was effective in reducing production of proinflammatory cytokines induced by IFN-α/β (fig. S5D) (*12*). Importantly, αIFNAR treatment from day 0 or day 3 post infection, had no effect on lung epithelial cell proliferation (fig. S5D). We therefore conclude that in our model of influenza virus infection in B6 mice, IFN-λ responses are most effective in disrupting epithelial regeneration during influenza recovery, and these effects are mediated directly in epithelial cells.

### Type I and III IFNs block AEC proliferation and differentiation

To understand mechanistically how IFNs exert the antiproliferative effects observed *in vivo*, we set up primary murine AEC cultures. Progenitor basal cells from distal airway sites are involved in the regeneration of the lung epithelium following influenza virus-induced damage (*6, 8, 30*). AEC cultures undergo rapid proliferation, and also differentiation upon exposure to an air-liquid interface (ALI), recapitulating repair processes observed in the lung and airway epithelium following damage *in vivo* (*31, 32*). Similar to IFNs used *in vivo*, IFNs used for *in vitro* assays were titrated on AEC cultures, to compare IFN subtypes at equivalent biological potency (fig. S6A and B). AEC cultures seeded at a low density in the presence of IFN-β or -λ were unable to reach confluence, whilst IFN-α treatment significantly reduced AEC number and the time taken for cultures to reach confluence (Fig. 2A left, and fig. S7A).

**Fig. 2.**
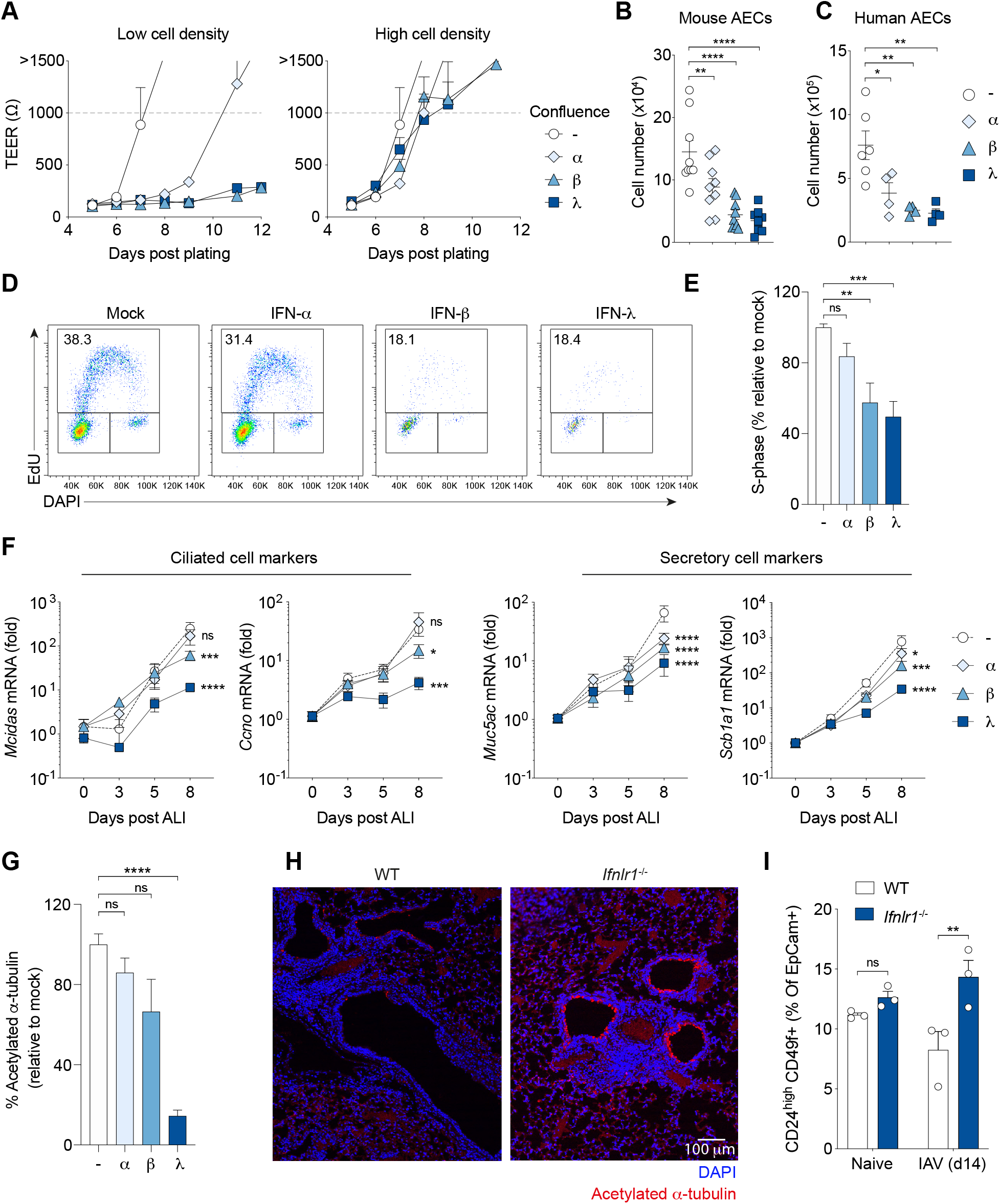
IFN signalling blocks airway epithelial cell growth and differentiation. (**A**) AECs were seeded at a low density (500 cells/transwell) or high density (10^4^ cells/transwell) in the presence of equivalent doses of IFN-α, -β, -λ, or media control, and grown for 12 days (n = 3 for all conditions). Confluence was determined by measuring transepithelial electrical resistance (TEER; >1000Ω = confluent cultures). (**B, D** and **E**) Proliferating mouse AEC cultures (2 days prior to exposure to an ALI) were treated for 5 days with IFNs (day −2 ALI to day 3 post ALI), and effects on growth were determined by cell number (B) (n = 9) and EdU incorporation to measure proliferation (D and E) (n = 9). (**C**) Primary human AEC cultures were treated with IFNs for 5 days, and cells were counted (n = 4 to 6). (**F** and **G**) Mouse AECs were grown to confluence, then exposed to an air-liquid interface (ALI) for 2 days. IFNs were then administrated for 6 days during ALI exposure (n = 6 for all conditions). Differentiation was determined by mRNA expression of the indicated genes (F) and the level of acetylated α-tubulin staining in cultures (G). (**H** and **I**) WT and *Ifnlr1*^-/-^ mice were infected with influenza virus, and lungs were analysed by immunofluorescence (DAPI/acetylated α-tubulin) on day 10 post infection (n = 4) (H), and flow cytometry (EpCam^+^CD49f^high^CD24^+^) on day 14 post infection (n = 3) (I). All data are representative of at least three independent experiments. Data are shown as mean ± s.e.m. and statistical significance was assessed by one-way (B, C, E and G) or two-way (F and I) ANOVA with Dunnett’s post-test. ns, not significant (P >0.05); \**P* ≤ 0.05, \*\**P* ≤ 0.01, \*\*\**P* ≤ 0.001, \*\*\*\**P* ≤ 0.0001.

To study growth inhibitory effects of IFNs without blocking the establishment of AEC cultures, we seeded cells at a higher density before treating with IFN for 5 days. All 3 IFN subtypes delayed the time taken for cultures to reach confluency, and impaired cell growth, with IFN-β and IFN-λ having the most significant effects (Fig. 2A right, and 2B). Growth inhibitory effects are dependent on the presence of the respective IFN receptor (fig. S7B). Similar effects were observed when primary human AEC cultures were treated with equivalent doses of IFN types (fig. S6C and D), with human IFN-β and -λ treatment resulting in the most significant decrease in cell number (Fig. 2C).

To gain mechanistic insight into IFN-mediated disruption of AEC growth, we analysed IFN treated AEC cultures by flow cytometry to study effects on apoptosis and proliferation. Consistent with the observed reduction in cell number, IFN-β or -λ treatment resulted in a significant increase in the frequency of apoptotic or necrotic cells (Annexin V^+^/TO-PRO-3^+^) (fig. S7C and D). We also observed a more than 50% reduction in the number of actively proliferating cells, as measured by incorporation of the thymidine analogue EdU, while IFN-α treatment had only a modest effect (Fig. 2D and E). These findings were confirmed by measuring CFSE dilution (fig. S7E and F, left). Interestingly, IFN-mediated antiproliferative effects were only observed in highly proliferative AEC cultures, as treating established cultures had no effect (fig. S7E to G, right). These data suggest that IFNs actively disrupt cell cycle progression without killing quiescent cells. Therefore, the increase in apoptosis observed (fig. S7C and D) may occur as a result of failed progression through the cell cycle following IFN treatment, as seen previously (*33*).

We next sought to examine the effects of IFNs on AEC differentiation. Following acute damage, populations of basal cells and Scgb1a1^+^ secretory cells give rise to secretory and multiciliated cell subtypes (*34*). Timely repopulation of airway epithelial cell subsets following damage is crucial in maintaining barrier function. Indeed, influenza virus infection has previously been shown to decrease mucociliary clearance in mice, resulting in an increased susceptibility to superinfection with *S. pneumoniae* (*35*). To study specific effects of IFNs on AEC differentiation, we initiated IFN treatment later during the course of AEC growth throughout exposure to an ALI, the time in which AEC differentiation is induced (fig. S8A). Similar to our findings for proliferation, IFN-β and -λ treatment significantly reduced the expression of genes pertaining to multiciliated (*Mcidas/Ccno*) and secretory cell (*Muc5AC, Scb1a1*) differentiation, with IFN-λ having the strongest effect (Fig. 2F). Expression of the basal cell marker *Krt5* remained unchanged, or increased by IFN-λ treatment, suggesting maintenance of stemness (fig. S8B). We also observed a significant reduction in the number of ciliated cells in AEC cultures (acetylated α-tubulin^+^) following IFN-λ treatment, but not with IFN-α or -β (Fig. 2G and fig. S8C). To confirm these findings *in vivo*, WT and *Ifnlr1*^-/-^ mice were infected with influenza virus, and lung sections were analysed by immunofluorescence. *Ifnlr1*^-/-^ mice had increased acetylated α-tubulin^+^ ciliated cells in repairing conducting airways on day 10 post infection (Fig. 2H). We quantified these findings by flow cytometry, and observed an increase in the frequency of differentiated AECs (EpCam^high^CD49f^high^CD24^+^) composed of ciliated, goblet and club cells (Fig. 2I and fig. S2)(*5*). From these results, we conclude that IFN-λ signalling reduces the capacity for basal cell differentiation during recovery from influenza virus infection.

### IFNs disrupt AEC growth via induction of p53-dependent downstream pathways

To understand mechanistically how IFNs mediate antiproliferative effects, we performed RNA-sequencing (RNA-seq) on IFN-treated AEC cultures (Fig. 3A). Principal component analysis (PCA) showed that untreated AEC control samples clustered depending on length of culture growth (including ALI exposure) along the principal component 2 (PC2), indicating that this axis represents proliferation and differentiation (Fig. 3B). AECs treated with IFN subtypes for 4 hours of culture clustered away from their respective mock on PC1, suggesting this principal component of variance (46%) is attributable to induction of antiviral ISGs (Fig. 3B). Additionally, 4-hour IFN treatment clustered samples together regardless of subtype (Fig. 3B), confirming the equal subtype dosage based on previous titrations (fig. S6A, B, and S9A).

**Fig. 3.**
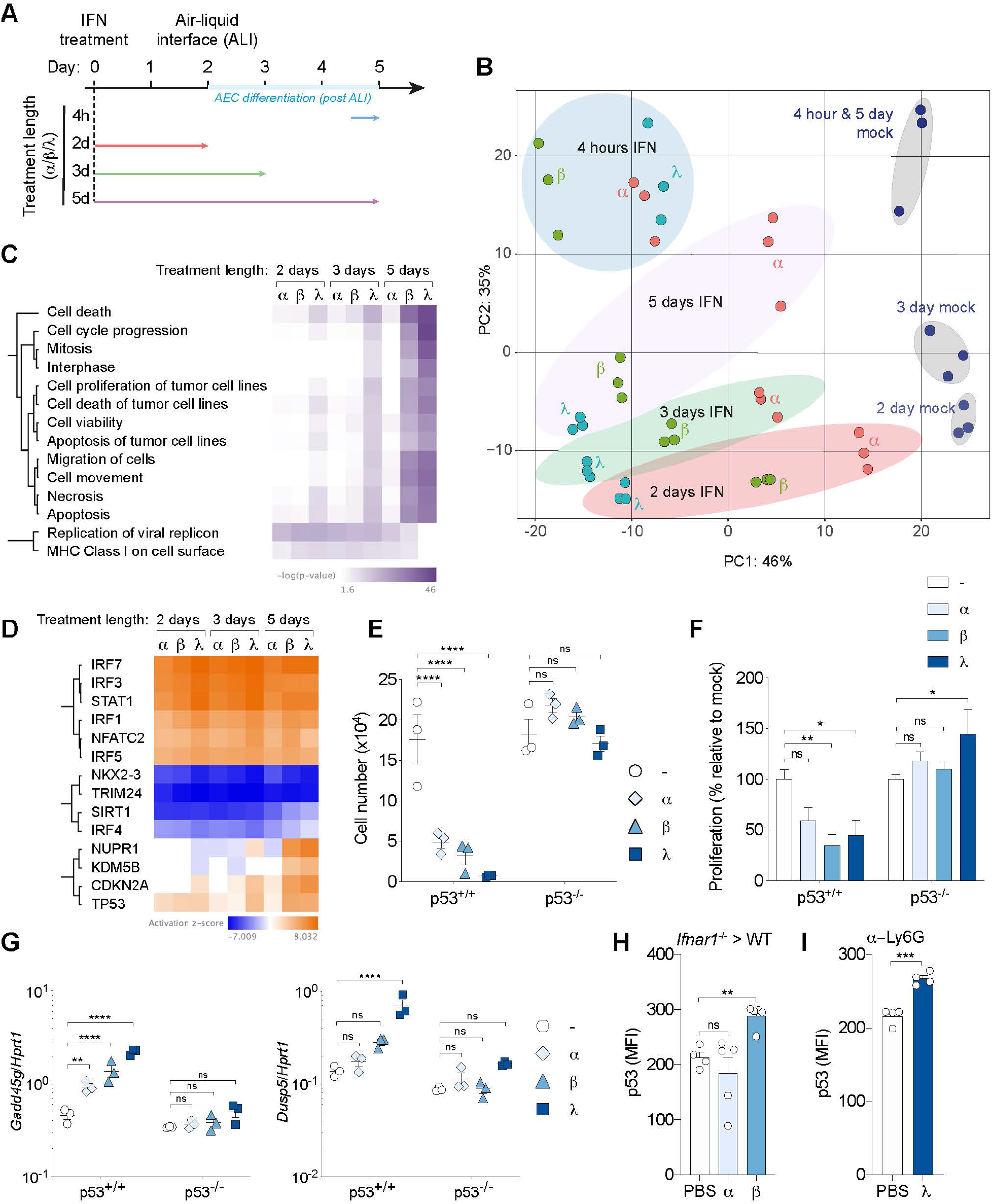
Type I and III IFNs activate antiproliferative and cell death pathways in AECs via induction of p53. (**A**) Schematic diagram for IFN treatment of AECs for RNA-seq analysis. (**B**) PCA plot of RNA-seq data from AECs following IFN treatment, and untreated controls. (**C**)Heatmap for significant differences in ‘Canonical Pathways’ for 9 pairwise comparisons between indicated IFN treatment and the respective mock, at each time point (fold change > 1.5, one-way ANOVA with Benjamini-Hochberg correction, P < 0.05), Gene expression was compared using IPA Comparison Analysis. (**D**) Predicted upstream transcriptional regulators of differentially expressed genes (IPA analysis). (**E** to **G**) WT and *p53*^-/-^ AECs were treated with IFN subtypes for 5 days, and measured for growth by cell number (E), CFSE dilution (F), and mRNA expression of indicated genes (G) (n = 3 for all conditions). (**H** and **I**) *Ifnarl*^-/-^ > WT BM chimeric mice (H) (n = 4 to 5) and α-Ly6G treated mice (I) (n = 4) infected with influenza virus (X31), and treated with IFN every 24 hours consecutively for 4 days (day 7 to 10 post infection), before EpCam^+^MHCII^+^CD49f^low^ AT2 cells were analysed for p53 mean fluorescence intensity (MFI) on day 11 post infection by flow cytometry. All data are representative of at least two independent experiments (E to I). Data are shown as mean ± s.e.m. and statistical significance was assessed by two-way (E to G) or one-way (H) ANOVA with Dunnett’s post-test, or unpaired two-tailed Student’s t-test (I). ns, not significant (P >0.05); \**P* ≤ 0.05, \*\**P* ≤ 0.01, \*\*\**P* ≤ 0.001, \*\*\*\**P* ≤ 0.0001.

AECs were also treated for 2, 3, or 5 days to understand the effects of prolonged IFN treatment in growing AEC cultures. 4-hour and 5-day IFN treated cultures shared the same experiment endpoint, and therefore share the same untreated controls (Fig. 3A). 5 days of IFN-β or -λ treatment, resulted in the clustering of samples away from corresponding untreated controls on both PC1 and PC2, indicating maintained ISG induction (PC1) and a block of AEC proliferation and development (PC2) (Fig. 3B). Conversely, the response induced by IFN-α was significantly reduced after just 2 days of treatment, consistent with previous findings (*36*). Ingenuity pathway analysis (IPA) of differentially expressed genes following prolonged IFN treatment (2, 3 and 5 days) revealed the top cluster pertaining to biological pathways regulating cell cycle and cell death, most significantly induced by IFN-λ treatment across all timepoints (Fig. 3C). Predicted upstream transcriptional regulators identified typical regulators of IFN function, including interferon regulatory factors (IRFs) and signal transducer and activator of transcription (STAT) proteins, in addition to regulators of cell cycle (Fig. 3D). We identified the tumour suppressor protein p53 as a top candidate regulating IFN-inducible antiproliferative effects, owing to its significant predicted activity following prolonged IFN-β and -λ treatment in AECs (Fig. 3D). Additionally, gene set enrichment analysis (GSEA) revealed IFN-mediated induction of the p53 downstream pathway and p53-regulated downstream targets (fig. S9B and C).

To confirm the role of p53, we utilised *p53*^-/-^ AEC cultures. IFN treatment had no effect on *p53*^-/-^ AEC number or proliferation following IFN treatment (Fig. 3E and F). IFN-mediated induction of antiproliferative downstream p53 target genes *Gadd45g* and *Dusp5* (*37, 38*) was p53 dependent, as IFNs had no such effect in *p53*^-/-^ AEC cultures (Fig. 3G). Additionally, IFN-λ had a much-reduced negative effect on multiciliogenesis in differentiating *p53*^-/-^ cultures, compared to its effect on WT and *p53*^+/-^ cultures (fig. S9D). We next sought to examine whether IFNs regulate p53 activity in epithelial cells during lung repair *in vivo*. To study IFN effects specifically in the lung epithelium, we once again generated *Ifnar1*^-/-^> WT bone marrow chimeric mice for IFN-α or –β treatment, or depleted neutrophils in WT mice with αLy6G for IFN-λ treatment (fig. S3A). In agreement with *in vitro* observations, IFN-β and -λ, but not IFN-α, significantly upregulated p53 expression in repairing lung epithelial cells (Fig. 3H and I). Taken together, these results demonstrate that IFN-β and -λ mediate antiproliferative effects in AECs via induction of p53. Additionally, our RNA-seq analyses confirm the potent antiproliferative activity of IFN-λ, compared to type I IFN subtypes.

### *Ifnlr1*^-/-^ mice have improved epithelial repair and barrier function following influenza virus infection

Our data supports a significant role for IFN signalling, in particular IFN-λ, in the reduction of epithelial proliferation and differentiation during lung repair. We therefore determined whether this alters the state of lung epithelia or affects barrier function of the respiratory epithelium. RNA-seq analysis of sorted influenza virus infected lung epithelial cells (EpCam^+^CD31^-^CD45^-^) from WT or *Ifnlr1*^-/-^ mice confirmed an upregulation of pathways pertaining to proliferation and multiciliogenesis in *Ifnlr1*^-/-^ mice (Fig. 4A). Improved repair correlated with reduced lung damage, with a reduction in both the total, and red blood cell number in the BALF of *Ifnlr1*^-/-^ mice day 8 post infection (Fig. 4B and C). Additionally, *Ifnlr1*^-/-^ mice had less inflammatory immune cell infiltrate dispersed in lung tissues (Fig. 4D). Influenza virus infections in humans are often exacerbated by opportunistic bacterial pathogen, including *S. pneumoniae* (*39*). A contributing factor driving this synergy, is a compromised epithelial barrier as a result of influenza virus-induced damage. To measure the effects of IFN-λ signalling on lung barrier function, we challenged mice with *S. pneumoniae* 8 days post influenza virus infection. We used BM chimeric mice to study specific effects of IFN-λ in epithelial cells. Mice lacking IFNLR in the stromal compartment (WT > *Ifnlr1*^-/-^) had improved survival following bacterial superinfection (Fig. 4E). Overall, our data indicate that IFN-λ signalling reduces the capacity for epithelial repair following influenza virus infection, which results in prolonged lung damage and compromised barrier function, consequently increasing susceptibility to bacterial superinfection.

**Fig. 4.**
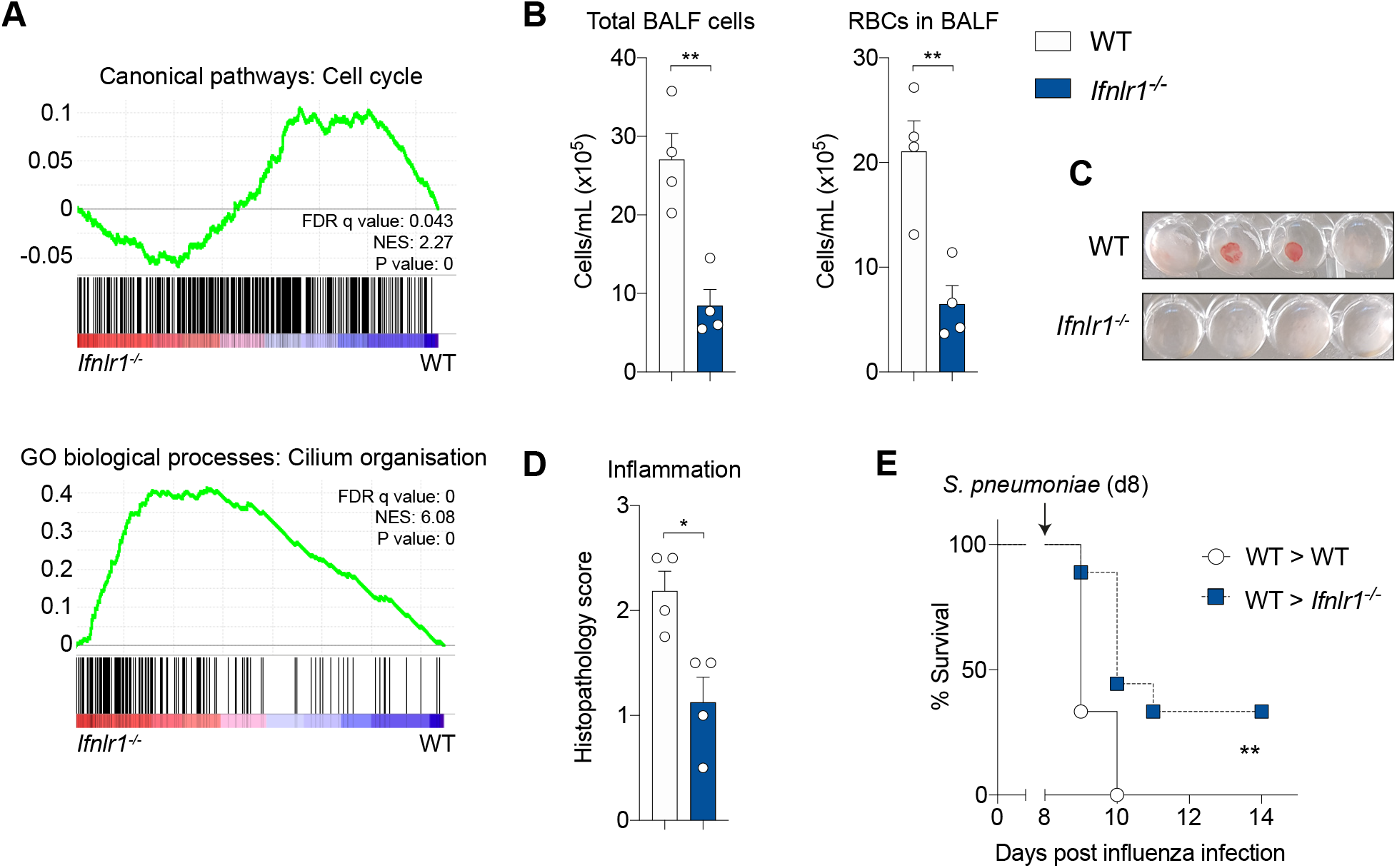
*Ifnlr1*^-/-^ mice have improved lung repair, reduced damage, and improved epithelial barrier function. WT or *Ifnlr1*^-/-^ mice were infected with 10^4^ TCID_50_ X31 influenza virus (X31). (**A**) GSEA plots of RNA-seq datasets from WT or *Ifnlr1*^-/-^ bulk lung epithelial cells (EpCam^+^) on day 8 post infection. (**B** and **C**) Total cell and red blood cell number in BALF on day 8 post infection (n = 4 for both WT and *Ifnlr1*^-/-^ mice). (**D**) Histopathological analysis of H&E lung sections on day 9 post infection (n = 4 for both WT and *Ifnlr1*^-/-^ mice). (**E**) Lethally irradiated WT and *Ifnlr1*^-/-^ mice were injected with WT bone marrow cells. Following reconstitution, chimeric mice were challenged with 2×10^5^ colony-forming units (c.f.u.) TIGR4 in 30μl on day 8 post influenza virus infection (n = 8 WT, n = 9 *Ifnlr1*^-/-^). All data are representative of at least two independent experiments (B to E). Data are shown as mean ± s.e.m. and statistical significance was assessed by unpaired two-tailed Student’s t-test (B to D) or log-rank (Mantel–Cox) test (E, survival). \**P* ≤ 0.05, \*\**P* ≤ 0.01.

## Discussion

Here we describe a novel mechanism by which type I and III IFN signalling aggravates lung pathology during respiratory viral infection. By focusing on the importance of epithelial repair during the recovery phase of infection, we highlight the potential for both IFN-α/β and IFN-λ in exacerbating influenza disease.

Our data indicates that IFN-λ is most effective in disrupting lung epithelial repair. Specific blockade of type I IFN responses with αIFNAR-treatment had no effect on lung epithelial proliferation. Comparison of IFN subtypes at equal biological potency in AECs revealed IFN-β and -λ to most significantly disrupt growth and induce antiproliferative pathways in primary human and murine epithelial cell cultures. Factors that determine the antiproliferative differential of IFN subtypes include: ligand-receptor binding affinity, length of stimulation, receptor density, and negative feedback control (*40–45*). Additionally, we found the magnitude of IFN-λ production in BALF was greater compared to IFN-α/β, extending into the crucial time window in which epithelial repair manifests post influenza virus infection. We therefore conclude that the reduced antiproliferative potential of IFN-α, combined with low IFN-α/β levels *in vivo*, limits the capacity for type I IFNs to block lung repair directly in B6 mice. However, we do not exclude a potential for host IFN-α/β production in directly blocking lung epithelial repair. This may be the case in situations of excessive type I IFN production and lung pathology, as seen in other mouse strains following influenza virus or SARS-CoV infection (*12, 13*).

Mechanistically, we identified the tumour suppressor protein p53 as the downstream regulator of IFN-induced antiproliferative effects. Research defining the mechanisms by which IFNs induce growth inhibitory effects have focused on type I IFNs, using transformed cell lines to explore antitumour therapeutics, with little understanding for their roles in normal physiology or during infection (*22*). In primary AECs, we observed no difference between WT and *p53*^-/-^ cultures in baseline growth or expression of downstream effector genes (including *Gadd45g* and *Dusp5*), indicating that p53 is inactive in these cells under normal conditions. Once activated, p53 triggers cell cycle arrest and apoptosis through induction of its target genes (*46*), and has previously been shown to directly regulate IFN-αβ antitumour and antiviral responses (*47*).

We also show that IFN-λ disrupts AEC differentiation during lung repair. The differentiation of AEC cultures triggered by air exposure is accompanied by reduced cell proliferation. As we treated cultures specifically during this period, the block in differentiation appears to be independent of antiproliferative IFN effects. Multiciliogenesis in AECs requires tightly coordinated use of the mitotic machinery for centriole amplification, required for the generation of cilia (*48*). We therefore speculate that IFN-induced p53 activity disrupts this coordination, thereby impairing AEC differentiation. Respiratory virus infections are associated with chronic lung disease such as asthma and chronic obstructive pulmonary disease (*49*), and proinflammatory cytokines have been shown to induce epithelial remodelling in human AECs (*50*). Therefore, our data suggest a mechanistic link between antiviral immune responses and longterm changes in lung physiology.

Pathogenic respiratory viruses, including IAV and CoV, are a significant global health concern owing to their potential to cause pandemics. The ongoing outbreak of CoV disease 2019 (COVID-19) presents the most severe global health crisis since the Spanish flu pandemic of 1918 (*51–53*). Similar to influenza, severe cases of COVID-19 are associated with massive alveolar damage and progressive respiratory failure (*54*), and SARS-CoV-2 has been shown to induce expression type I and III IFNs in primary human AEC cultures (*55*). In addition, histopathological findings in post-mortem lung biopsies of COVID-19 victims reveal neutrophilic infiltration consistent with superimposed bacterial bronchopneumonia (*56*). Causes of variation in host disease severity during respiratory virus infection are not well understood. Studies comparing influenza patient cohorts found little or no correlation between disease severity and viral load (*57–60*), suggesting a role for genetically determined host factors, including mechanisms amplifying immunopathology.

Attempts to gain a better understanding of human influenza pathogenesis have largely been restricted to whole blood expression analysis (*61–63*), and therefore exclude localised immune and epithelial repair responses in lung tissues (*64*). However, macaques infected with the highly virulent 1918 H1N1 strain, revealed an elevated ISG signature late during infection bronchial tissue (*65*). Additionally, SARS-CoV patients displayed robust IFN responses, and aberrant ISG expression in severe cases (*66*). Whether susceptible individuals display excessive IFN responses late during infection, and the potential impact this has on lung repair, remains to be determined. Our transcriptomic analyses of regenerating epithelial cells *in vivo* link host IFN-λ responses with a reduction in the capacity for lung repair. Analysis of lung tissue and BALF from respiratory virus infected patients experiencing severe disease will provide insight into the mechanisms regulating disease pathogenesis, including potential interplay between IFNs and epithelial cells.

Lack of an available vaccine during the emergence of novel zoonotic viruses, like SARS-CoV-2, highlights the importance of generating and implementing effective host-directed antiviral and immunomodulatory treatments. Our data highlights the importance of a well-timed IFN response, with regards to both host immunity and exogenous treatment. IFN-α/β treatment exacerbates inflammation and disease severity in mice if administered past the peak of viral replication during influenza virus or Middle East respiratory-syndrome-CoV (MERS-CoV) infection (*67, 68*). IFN-λ treatment early during influenza virus infection was shown to be protective, owing to its ability to confer antiviral protection without generating the proinflammatory responses associated with IFN-α/β (*68, 69*). For these reasons, IFN-λ is therefore being implemented as a candidate therapy for COVID-19 (*70*). However, our findings here identify a mechanism by which IFN-λ exacerbates respiratory virus disease acting specifically through the respiratory epithelium, independent of immunomodulation.

Taken together, these data indicate the necessity for effective regulation of host IFN responses, and the importance of timing and duration when considering IFNs as therapeutic strategies to treat respiratory virus infections. Optimal protection would be achieved by strong induction of ISGs early during infection to curb viral replication, followed by timely downregulation of IFN responses, enabling efficient lung epithelial repair. These data change our understanding of immunopathology during viral infection, and have important implications for IFN usage as antiviral therapy.

## Supporting information

Major Supplemental Material

Major Supplemental data S1

## Acknowledgments

We thank the Crick sequencing, histopathology, flow cytometry and animal facilities for excellent support. We are grateful to Simon Priestnall and Alejandro Suarez-Bonnet for histopathology scoring of lung H&E sections, and to Anne O’Garra, Gitta Stockinger, Peter Staeheli and Daniel Schnepf for critically reading the manuscript.

## Author contributions

Conception: A.W., S.C. and J.M; research design and experimentation: J.M., S.C. and T.McC.; data analysis: J.M., S.C., T.McC. and M.L.; production and provision of key reagents: H.H.G. and R.H.; writing and editing of the manuscript: J.M., A.W., and S.C. All authors read and approved the final manuscript.

## Data and materials availability

All data supporting the findings of this study are available within the paper or in the supplementary materials, or are available from the authors upon request.

## Supplementary Materials

Materials and Methods

Figures S1-S9

Tables S1

Data S1

References (*31, 71–75*)

