## Supplementary material for "Type I and III interferons disrupt lung epithelial repair during recovery from viral infection": Major Supplemental Material

**This PDF file includes:**

Materials and Methods
Figs. S1 to S9
Tables S1
Caption for Data S1

**Other Supplementary Materials for this manuscript include the following:**

Data S1: Differential gene expression in 4-hour IFN treated versus mock AECs cultures.

### Materials and Methods

*Mice*

All experiments used male and female mice at 6–12 weeks of age bred at the Francis Crick Institute (2016–2020) under specific pathogen-free conditions. All animal experiments were approved by the Home Office, UK, under project license P9C468066, and carried out in accordance with the Animals (Scientific Procedures) Act 1986 and the GSK Policy on the Care, Welfare and Treatment of Animals. All genotypes were bred on a C57BL/6J background and maintained as homozygous lines. Genotypes were C57BL/6J, *Ifnar1*$^{-/-}$, *Ifnlr1*$^{-/-}$ (provided by A. O'Garra at the Francis Crick Institute), *p53*$^{-/-}$ (*in vitro only*), and B6.A2G-Mx1 congenic mice carrying functional Mx1 alleles on the C57BL/6 background (*71*) (a kind gift from Dr P. Staeheli, Freiburg Univ.).

To generate BM chimeras, WT, *Ifnar1*$^{-/-}$ and *Ifnlr1*$^{-/-}$ recipient mice were lethally irradiated (2 x 6.5 Gy, 15-hour interval) and reconstituted with 5 x $10^6$ donor BM cells from WT or *Ifnar1*$^{-/-}$ donor mice (as indicated in figures). BM chimeric mice were then left for 8 weeks before infection to reconstitute hematopoietic compartment.

*Influenza viruses*

X31 (a gift from J. Skehel, MRC-NIMR) and hvPR8-ΔNS1(1-126) (a gift from Dr P. Staeheli, Freiburg Univ. (*72*)) were grown in the allantoic cavity of 10 d-embryonated hen's eggs and were free of bacterial, mycoplasma and endotoxin contamination. All viruses were stored at −80 °C and titrated on Madin–Darby canine kidney cells. Virus was quantified in infected lungs by qPCR for the influenza *Matrix* gene on cDNA from whole lungs, normalized to the housekeeping gene *Hprt*.

*Infections*

C57BL/6J, *Ifnar1*$^{-/-}$ and *Ifnlr1*$^{-/-}$ mice were infected with X31 (10,000 $TCID_{50}$ in 30 µl PBS), and B6.A2G-Mx1 mice were infected with hvPR8-ΔNS1(1-126) (2,500 $TCID_{50}$ in 30 µl PBS). BM chimeric mice were infected with *S. pneumoniae* (TIGR4) (2x$10^5$ c.f.u. in 30uL PBS, day 8 post influenza infection. All infections were performed under light anesthesia (3% isoflurane) intranasally. With the exception of chimeras, mice were infected at 8 to 12 weeks of age. Chimeric mice were infected at 16 to 20 weeks of age. Pre-infection body weights were recorded and mice were weighed daily (at similar times of day) and monitored for clinical symptoms. Mice reached the clinical endpoint following the loss of 25% of their initial starting weight or at a clinical score of $>$ 5. Clinical scores were determined by (1 point each) piloerection, hunched posture, partially closed eyes, laboured breathing, hypothermia, decreased movement, movement only on provocation or (2 points) absence of movement on provocation or (5 points) middle-ear infection (disrupted balance).

*Mouse treatments*

*In vivo* IFN treatments in WT, B6.A2G-Mx1 and *Ifnar1*$^{-/-}$ > WT BM chimeric mice (subtype indicated in figure panels) were performed day 7 to day 10 post infection every 24 hours consecutively. IFNs were administered on day 7, 8 and 9 intraperitoneally (in 200 μl PBS) and on day 10 intranasally (in 50 μl PBS under light anesthesia (3% isoflurane). IFNs were used at equal biological potency based on titrations: IFN-α 8 μg/dose (Universal IFN-α, kind gift from Dr P. Staeheli, Freiburg Univ.); IFN-β $10^6$ U/dose (PBL); IFN-λ 2.5 μg/dose (recombinant mouse IFN-λ2) (*73*). WT mice were treated with α-Ly6G monoclonal antibody (clone 1A8) or Isotype control (150 μg per 200 μl intraperitoneally) every 24 hours, between day 5 and 10 post infection (IgG2b). WT mice were treated with α-IFNAR (MAR1-5A3, Bio X Cell) (400 μg per 200 μl intraperitoneally) on the day of infection (day 0) or from day 3 post infection, every 48 hours.

*Primary AEC cultures and in vitro IFN treatments*

Isolation and culture of primary murine AECs were performed as previously described. In brief, tracheal cells isolated by enzymatic treatment were expanded in a T-75 flask to 100% confluence with Rho kinase inhibitor Y27632 10uM (*74*). Cells were then trypsinised and seeded ($10^4$ cells/transwell, unless otherwise indicated) 0.4 μm pore size clear polyester membrane (Greiner) coated with a collagen solution. Cells were grown in submersion until confluent, and then exposed to air to establish ALI. Human biological samples were sourced ethically and their research use was in accord with the terms of the informed consents. Primary human bronchial epithelial cells were purchased from Lonza and cultured as per manufacturer's instructions. In brief, cells were expanded in a T-75 flask to 60% confluence and then harvested for seeding onto transwells at $5x10^4$ cells per insert. At confluence, liquid was removed from the upper chamber to establish ALI.

Murine and human AECs were treated with IFN 1-day post plating (2 days prior to confluency and therefore ALI exposure) for 5 consecutive days (for proliferation assays), or 3 days post plating (2 days post confluency/ALI exposure) for 6 consecutive days (to study effects on differentiation). Concentration of mouse IFNs based on titrations for equal biological potency were as follows: 600 U/ml IFN-$\alpha_4$ (PBL), 1000 U/ml IFN-β (PBL), 5ng/ml for both IFN-λ2 and -λ3 (R&D). Concentration of human IFNs based on titrations for equal biological potency were as follows: 1ng/mL Universal IFN-α, (Dr P. Staeheli), 2.3ng/ml h-IFN-β (PBL), 3.3ng/ml h-IFN-λ3 (R&D) (*74*).

*Flow cytometry*

For cell isolation from lung tissues, mice were euthanised (600mg kg$^{-1}$ pentobarbital + 17mg kg$^{-1}$) mepivacaine and then perfused with 10 ml of ice-cold PBS through the right ventricle of the heart. 1.5ml Dispase II (5 mg/ml in AB-IMDM) (Sigma) was then injected intratracheally into the lungs, followed by 0.4 ml 1% low-gelling agarose solution (in PBS) (Sigma). Mice were then placed on ice allowing the agarose/dispase-filled lungs to set. Lungs were then dissected, and placed in 2 ml Dispase II solution for 30 minutes to dissociate epithelial cells. Lungs were passed through a 100 µm filter, before a 10-minute DNase I digestion (50 µg ml$^{-1}$) (Sigma). Following digestion, lung homogenates were passed through a 70 µm filter, and centrifuged at 1,400 r.p.m. for 5 min at 4 °C, before red blood cell lysis. Single cell suspensions were preincubated blocked with anti-FcgRIII/II (Fc block), before a 30-min incubation with one or more fluorochrome-labelled antibodies. Cells were permeabilised with the Foxp3/Transcription factor staining kit (eBioscience) prior to intracellular staining for Ki67 or p53.

For EdU assays, AEC cultures were cultured with 2.5 µM EdU (Invitrogen) for 3 hours, before EdU$^{+}$ AECs were labelled using the Click-iT Plus EdU Alexa Fluor 488 Flow Cytometry Assay Kit (Invitrogen), according to the manufacturer's instructions.

Cells were labelled with CFSE (1 µM) (Molecular Probes) for 8 minutes, followed by 3x washes with media + sera. CFSE labelled AECs were trypsinized 24 hours post-labelling. Lung and AEC cells were analysed using a Fortessa X20 (Becton Dickinson).

*RNA isolation*

Lungs were harvested and stored in RNALater in -80$^{o}$C. Lungs were homogenised with a Kinematica Polytron PT 10-35 homogenizer in 3 ml RLT buffer (QIAGEN) + β-mercaptoethanol. AEC cultures were lysed in 350 µl RLT + β-mercaptoethanol. RNA was isolated using the Qiagen RNeasy mini kit, according to the manufacturer's instructions. 500 ng total RNA was reverse-transcribed using the qPCRBIO cDNA synthesis kit as per manufacturer's instructions. RT-qPCR was performed on an Applied Biosystems Quantstudio 3 RT-qPCR machine with 1x qPCRBIO Probe Mix Lo-ROX, and 1x Taqman primers. Results were normalized to the housekeeping gene *Hprt*.

The following probes (Applied Biosystems) have been used: *Hprt1* (Mm00446968_m1),
*Ccno* (Mm01297259_m1), *Mcidas* (Mm01308202_m1), *Muc5ac* (Mm01276718_m1), *Muc5b* (Mm00466391_m1), *Scb1a1* (Mm00442046_m1), *Dusp5* (Mm01266106_m1), *Gadd45g* (Mm0135255 0_g1), *ifi203* (mm00492601_m1), *Oasl2* (mm00496187_m1), *Rsad2*

(mm00491265_m1), *Rsad2* (human, Hs00369813_m1), *Mx1* (human, Hs00895608_m1), *Oas1* (human, Hs00973637_m1). Primers for influenza virus *Matrix* gene were as follows (*75*):

- forward: 5′-AAGACCAATCCTGTC ACCTCTGA-3′;
- reverse: 5′-CAAAGCGTCTACGCTGCAGTCC-3′;
- and probe: 5′-TTTGTGTTCACGCTCACCGT-3′.

*RNA-seq*

Following infection, epithelial cells were enriched by MACS separation ($CD45^-$) and cell sorting using a FACS Aria III (BD Biosciences) (Live single cells, $EpCam^+CD31^-$), and RNA was extracted. RNA-sequencing was performed on the HiSeq 4000 system (Illumina) with Single End 75 bp reads.  Read quality trimming and adaptor removal was carried out using Trimmomatic (version 0.36). Reads were aligned to the mouse genome (Ensembl GRCm38 release 89) using STAR (version 2.5.2a) and gene level counts were obtained using the RSEM package (version 1.2.31). For RSEM, all parameters were run as default except "-forward-prob" that was set to 0 for AECs (RN200717) and 0.5 for the lung samples (RN19197). Differential expression analysis was carried out with DESeq2 package (version 1.20.0) within R version 3.5.1. Genes were considered to be differential expressed with padj $\leq 0.05$.

Gene Set Enrichment analysis (GSEA, version 2.2.3) was performed for each pairwise comparison using gene lists ranked using the Wald statistic. Gene set pre-ranked analysis was carried out using C2 canonical pathways v5.2 and C5 biological processes v5.2. All parameters were kept as default except for enrichment statistic that was changed to classic and the max size which was changed to 500,000. Gene signatures were considered significant if FDR q-value % 0.05. IPA analysis was performed using differentially expressed genes with a fold change > 1.5 with $P < 0.05$.

*Histology and immunofluorescence*

Whole lungs were inflated with intratracheal injection of 4% paraformaldehyde (PFA), and fixed overnight also in 4% PFA, then embedded in paraffin and sectioned. Lung specimens were stained with haematoxylin and eosin (H&E) and then subjected to gross and microscopic pathologic analysis.

For immunofluorescent staining of paraformaldehyde-fixed AECs and dewaxed lung specimens, samples were permeabilized with 0.1% Triton X-100 for 15 min at room temperature and then blocked in 0.5% bovine serum albumin for 1 h. Primary antibodies specific for acetylated $\alpha$-tubulin (T7451, Sigma) were added apically in 0.5% bovine serum albumin and incubated for 1 h at room temperature. After washing with PBS, fluorochrome-conjugated secondary antibodies (Alexa Fluor, Life Technologies) were added for 1 h at room temperature. Finally, transwell filters were washed in PBS and mounted on slides using Vectashield mounting medium with 4,6-diamidino-2-phenylindole (Vector labs). Images were acquired on an Olympus VS120 slide scanner. Measurements of acetylated a-tubulin staining were made using regions of interest that cover the whole transwell. Binary images were generated by global thresholding and measured using the ImageJ 1.4j application.

*Quantification of IFNs and cells in BALF*

Mice were euthanised (600 mg $kg^{-1}$ pentobarbital + 16 mg $kg^{-1}$ mepivacaine) and BALF was collected through catheter insertion intratracheally, injecting and retracting 400-500 µl of PBS. BALF was centrifuged at 1,400 r.p.m. for 5 min at 4 °C, and supernatants were stored at −80 °C. Pelleted cells in the BALF were counted, and analysed by flow cytometry to determine the frequency of Ter-$119^{+}$ red blood cells. Concentrations of IFN-α/β were measured using 2-Plex ProcartaPlex (Invitrogen). IFN-λ (R&D) and IL-6 (eBioscience) were measured by ELISA, according to manufacturer's instructions.

*Statistical analysis*

Data shown as the means ± s.e.m. Sample sizes were designed to give statistical power, while minimizing animal use. All statistical comparisons were performed using Prism 8 (GraphPad). Figure legends denote the specific statistical tests used for each experiment. Statistical significance was determined as $P < 0.05$.

*Data availability*

All data supporting the findings of this study are available within the paper or in the supplementary materials, or are available from the authors upon request.

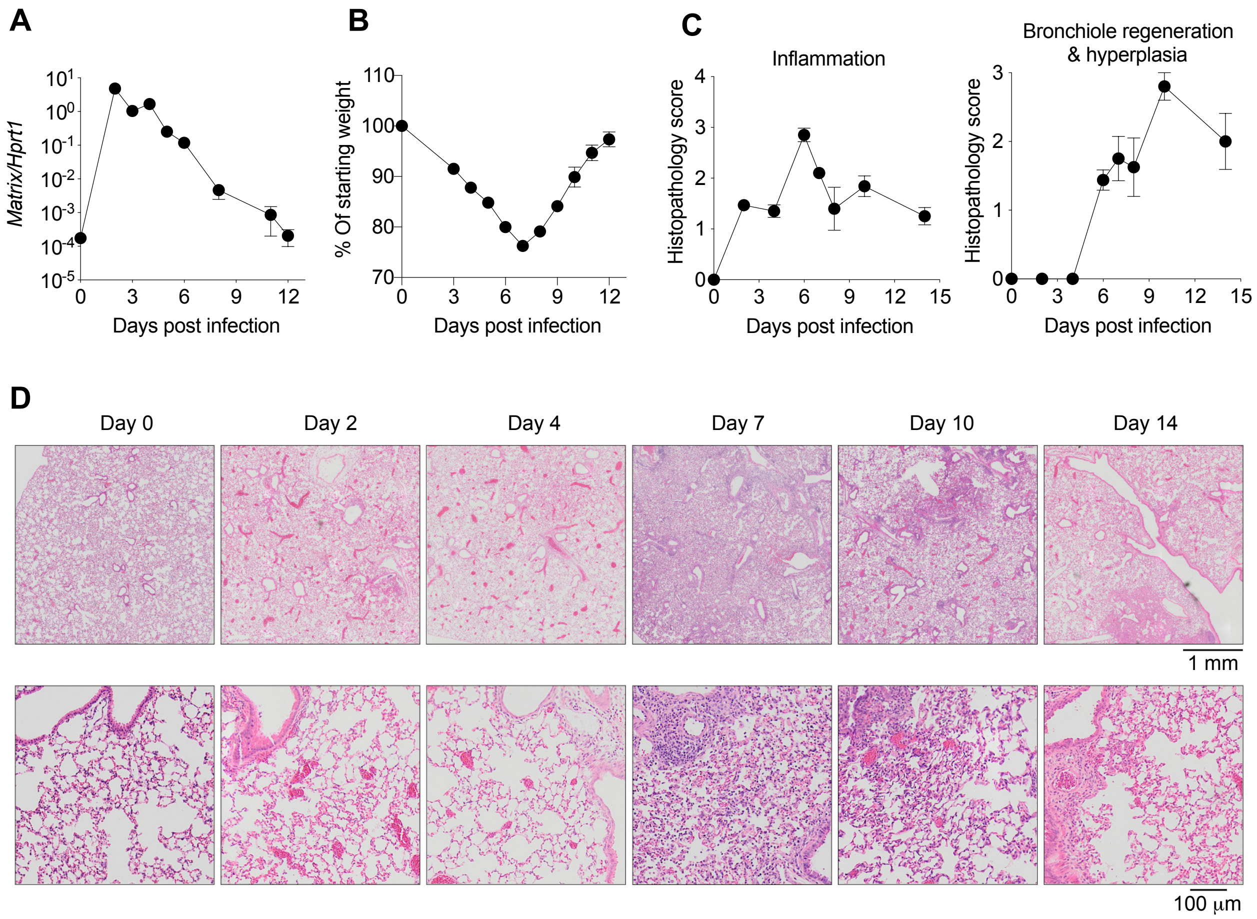


**Fig. S1. Dynamics of lung epithelial repair and IFN production during influenza infection.** WT mice were infected with 10,000 $TCID_{50}$ X31 (H3N2) influenza virus in 30 µl intranasally, and assessed for (**A**) influenza *M1* mRNA expression in total lung RNA (n = 6 mice per day), (**B**) weight loss (n = 5 mice per day), (**C** and **D**) and lung pathology (n = 5 mice per day). All data are representative of at least three independent experiments. Data are shown as mean ± s.e.m.

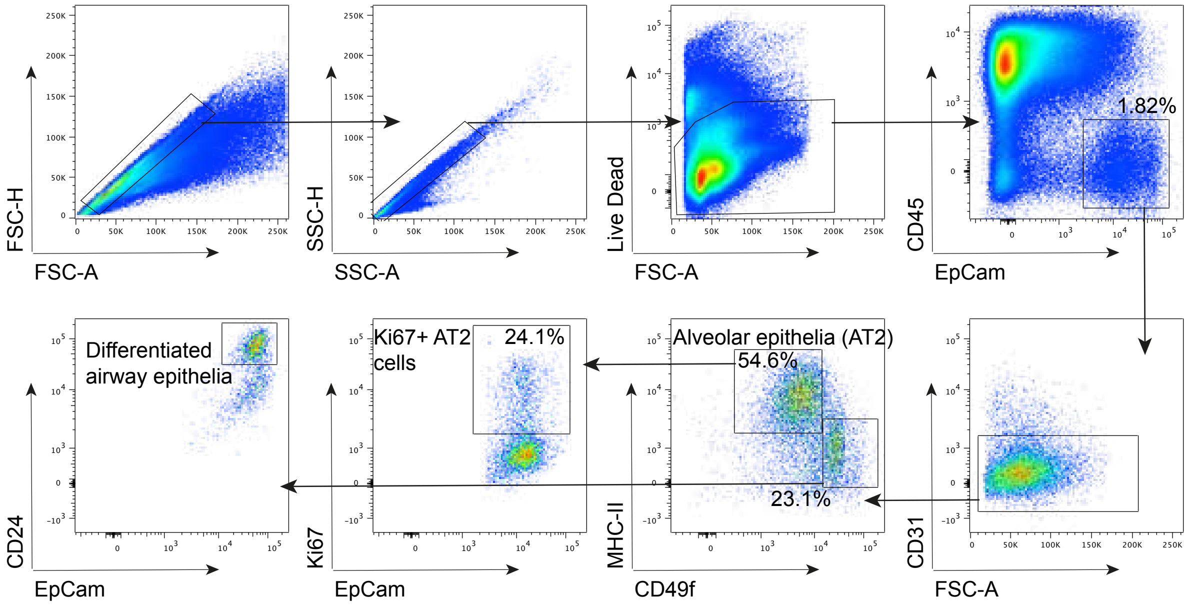


**Fig. S2. Lung epithelial cell analysis by flow cytometry.** Gating strategy for lung epithelial cells (WT, day 9 post influenza virus infection).

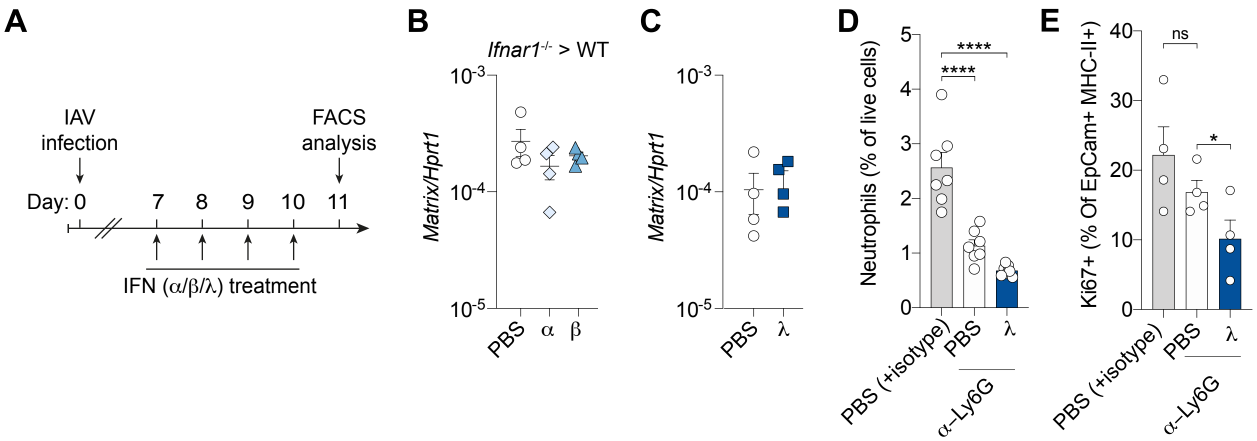


**Fig. S3. IFN treatment reduces lung epithelial cell proliferation independent of virus load and immune cells.** (**A**) Schematic for IFN treatment during lung regeneration phase of influenza virus infection. (**B** and **C**) Influenza virus matrix gene expression (M1) on day 11 post influenza virus infection in *Ifnar1*$^{-/-}$ > WT bone marrow chimeric mice (n = 4) (B) and WT mice (n = 4) (C). (**D**) Ly6C$^{hi}$CD11b$^{+}$Lineage (CD3/CD19/NK1.1/CD64/Siglec F)$^{-}$ neutrophil frequency following isotype control, or α-Ly6G treatment in influenza virus infected lungs on day 11 post infection. (**E**) Infected mice were treated with α-Ly6G monoclonal antibody (clone 1A8), or isotype control (n = 4) every 24 hours (day 5 to 11 post infection). α-Ly6G treated mice were also treated with IFN-λ (n = 4) or PBS control (n = 4), and proliferating (Ki67$^{+}$) AT2 cells (EpCam$^{+}$MHCII$^{+}$CD49f$^{low}$) were measured by flow cytometry on day 11 post infection. All data are representative of at least two independent experiments. Data are shown as mean ± s.e.m. and statistical significance was assessed by one-way ANOVA with Dunnett's post-test. ns, not significant (P >0.05); *$P \leq 0.05$, ****$P \leq 0.0001$.

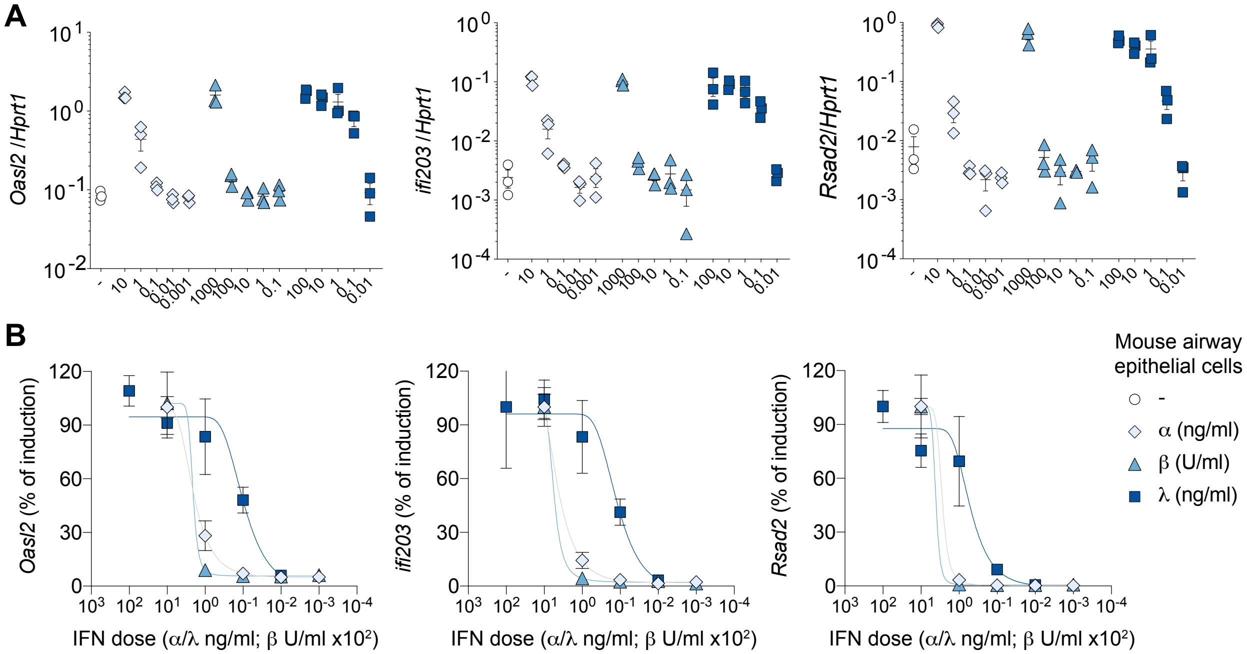


**Fig. S4. IFN titration for equal subtype dosage (IFNs used *in vivo*).** (**A**) Primary murine AEC cultures were treated for 4 hours at indicated concentrations with α, β and λ IFNs used in *in vivo* experiments. mRNA expression of indicated of antiviral ISGs was assessed by qPCR (n = 3 for all conditions) in IFN treated and untreated AECs. (**B**) Dose-response curves (Sigmoidal, 4PL, X is log(concentration)) were generated for each IFN subtype, relative to ISG expression at the highest IFN concentration.

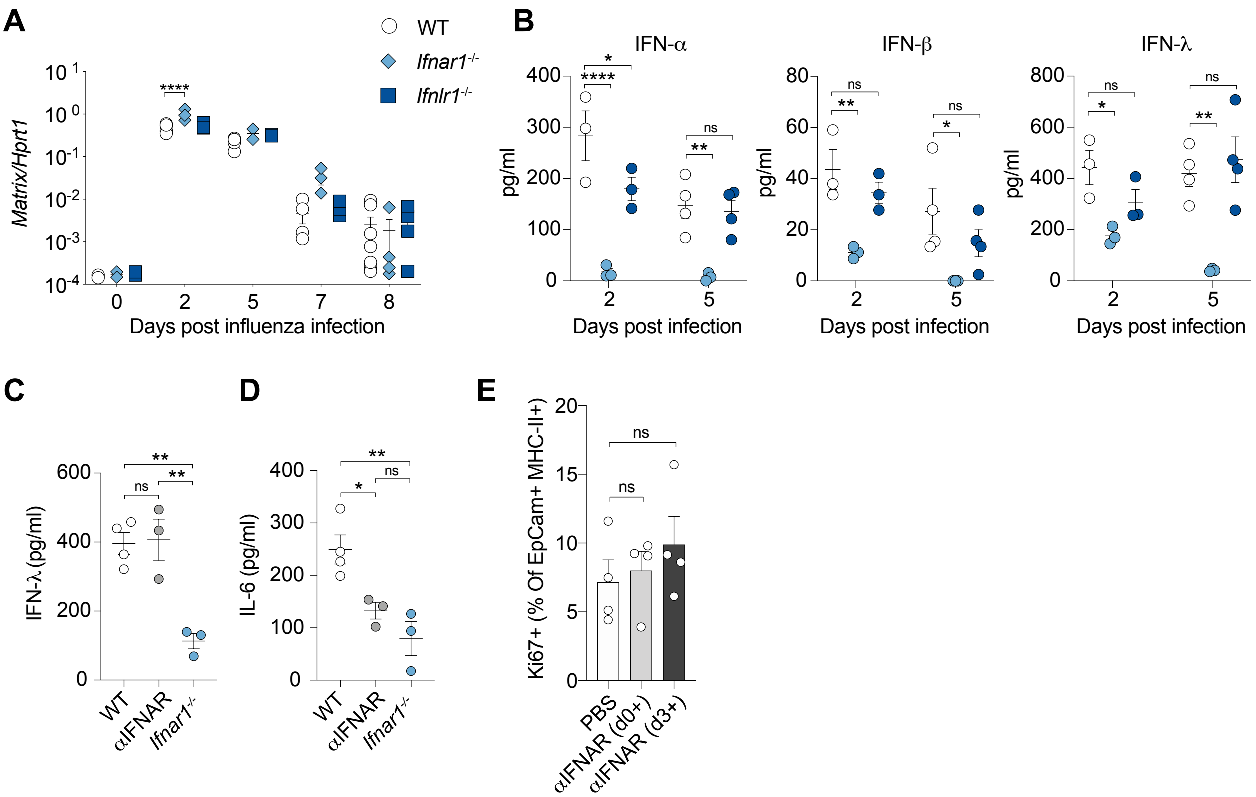


**Fig. S5. Improved lung epithelial cell proliferation is independent of changes in viral burden and type I IFN signalling, but dependent on IFN-λ signalling.** (**A** and **B**) WT (n = 3 to 5), *Ifnar1*$^{-/-}$ (n = 3 to 4), and *Ifnlr1*$^{-/-}$ (n = 3 to 4) mice were infected with influenza virus, and influenza virus matrix gene expression in the lung (A), or IFN-α, -β, and -λ levels in BALF (B) was measured at indicated days. (**C** and **D**) IFN-λ and IL-6 levels in BALF day 3 post infection in WT (n = 4) (n = 4), αIFNAR monoclonal antibody (MAR1-5A3) treated (day 0 and day 2 post infection) (n = 3), and *Ifnar1*$^{-/-}$ mice (n = 3). (**E**) Influenza virus infected WT mice were treated with PBS (n = 4) or αIFNAR monoclonal antibody (day 0, 3 and 5, or day 3 and 5) (n = 4 for both treatment groups), and Ki67$^{+}$ AT2 cells (EpCam$^{+}$MHCII$^{+}$CD49f$^{low}$) were detected by flow cytometry on day 8 post infection. All data are representative of at least two independent experiments. Data are shown as mean ± s.e.m. and statistical significance was assessed by two-way (A and B) or one-way (C to E) ANOVA with Tukey's (C and D) or Dunnett's (E) post-test. ns, not significant (P >0.05); *$P \leq 0.05$, **$P \leq 0.01$, ****$P \leq 0.0001$.

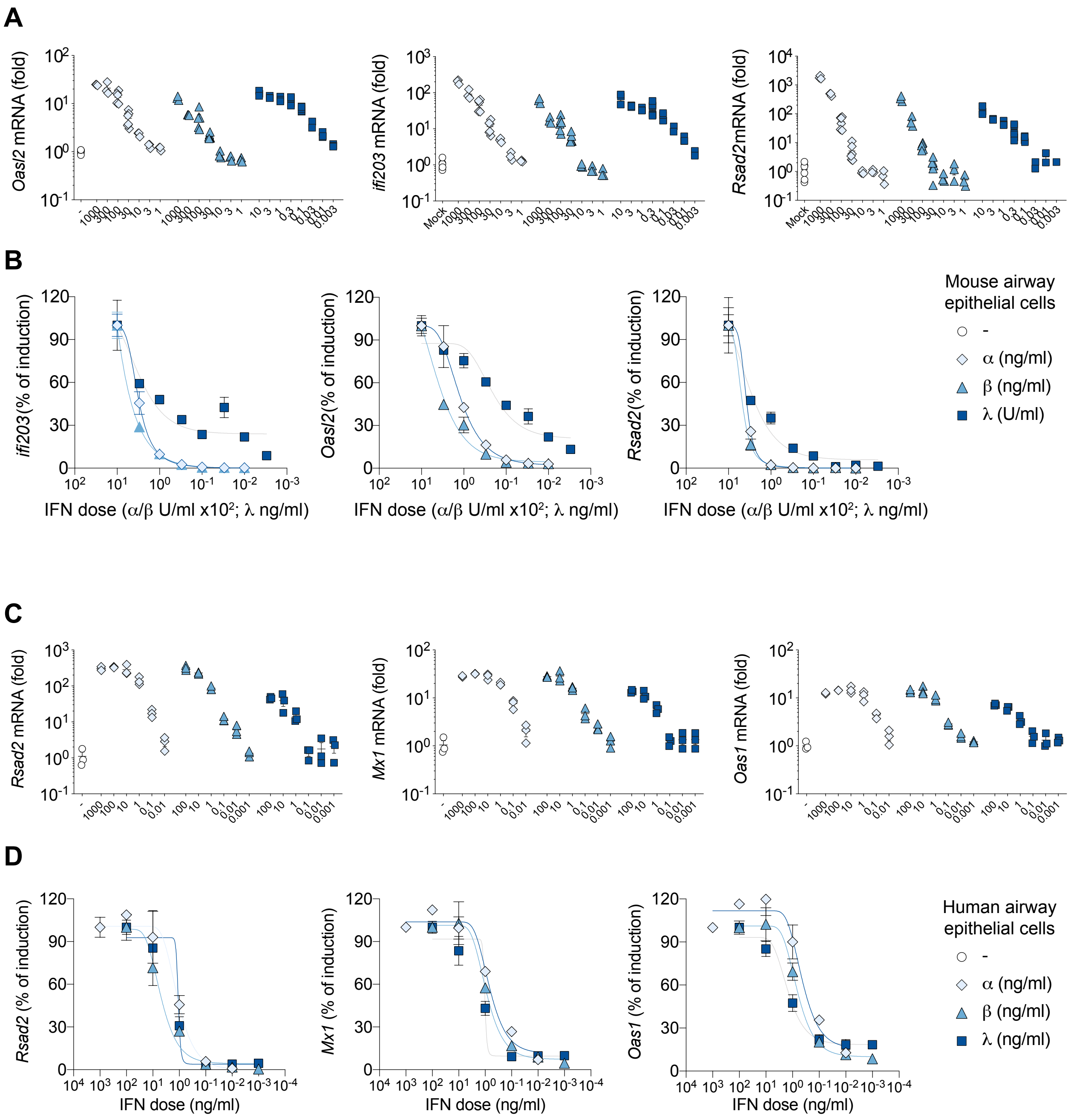


**Fig. S6. IFN titration for equal subtype dosage (IFNs used *in vitro*).** (**A** and **C**) Primary murine (A) and human (C) AEC cultures were treated for 4 hours at indicated concentrations with α, β and λ IFNs used in *in vitro* experiments. mRNA expression of indicated of antiviral ISGs was assessed by qPCR (n = 3 for all conditions) in IFN treated and untreated AECs. (**B** and **D**) Dose-response curves (Sigmoidal, 4PL, X is log(concentration)) were generated for each IFN subtype, relative to ISG expression at the highest IFN concentration.

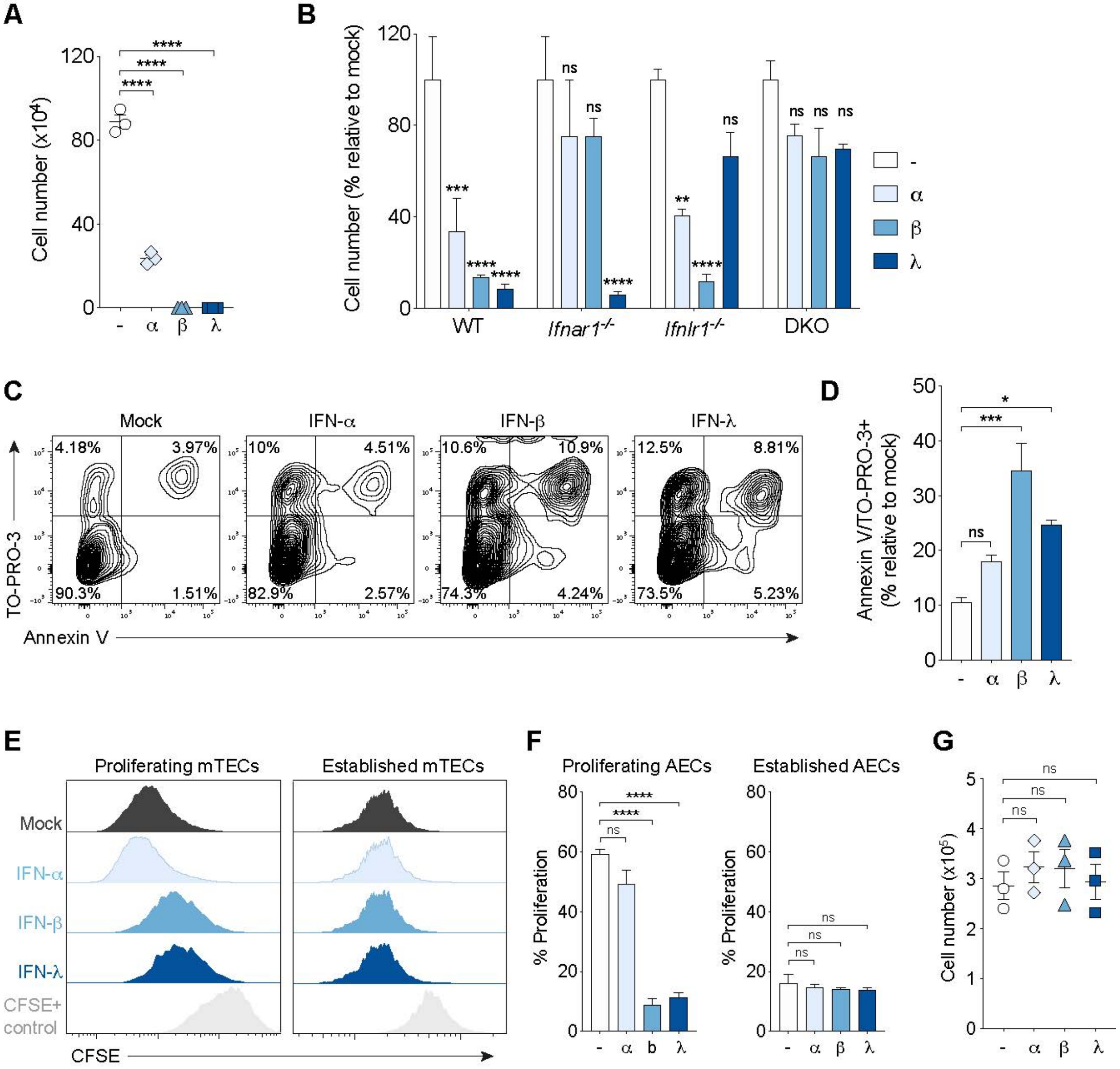


**Fig. S7. Dynamics for IFN-induced antiproliferative and proapoptotic effects in AECs.** (**A**) Cell number of WT AECs seeded at 500 cells/transwell in the presence of IFN-α, -β, -λ, or media control (n = 3 for all conditions). (**B**) Cell number of WT, *Ifnar1*$^{-/-}$, *Ifnlr1*$^{-/-}$, and *Ifnar1*$^{-/-}$*Ifnlr1*$^{-/-}$ (double receptor knockout) AEC cultures treated for 5 days with IFN subtypes or media control (n = 3 for all conditions). (**C** and **D**) AECs were treated for 5 days with IFN subtypes or media control, and the frequency of apoptotic or necrotic AECs (Annexin-V$^{+}$TO-PRO-3$^{+}$) were assessed by flow cytometry (n = 4 for all conditions). (**E** and **F**) Proliferating AEC (as in A to D, day -2 pre-exposure to an ALI) and established AEC (day 7 post-exposure to an ALI) cultures were treated for 5 days with IFN or media control. Cells were labelled with CFSE for the final 24 hours, and proliferation was measured by CFSE dilution (n = 3 for all conditions). (**G**) Established AEC cultures were treated for 5 days with IFNs or media control, and cells were counted (n = 3 for all conditions). All data are representative of at least four independent experiments. Data are shown as mean ± s.e.m. and statistical significance was assessed by one-way (A, D, F and G), two-way (B) ANOVA with Dunnett's post-test. ns, not significant ($P > 0.05$); *$P \leq 0.05$, **$P \leq 0.01$, ***$P \leq 0.001$, ****$P \leq 0.0001$.

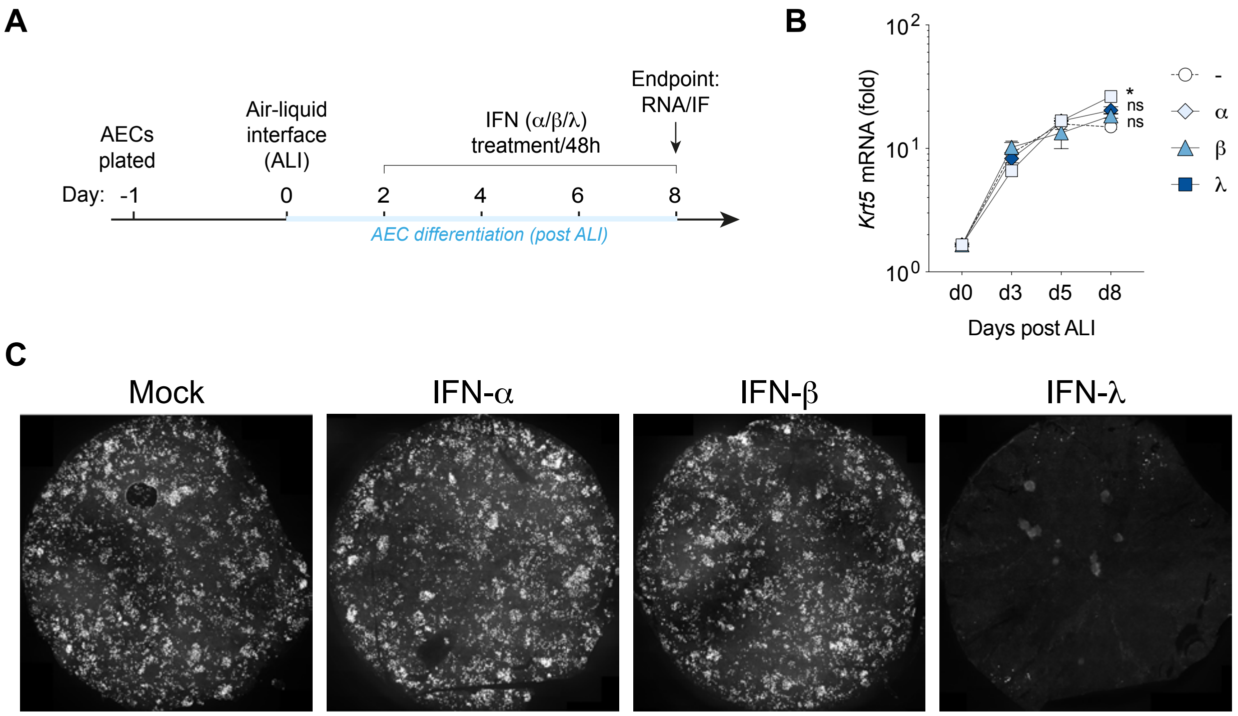


**Fig. S8. IFN-λ reduces AEC multiciliogenesis.** (**A**) Schematic diagram for IFN treatment of AEC cultures during differentiation (ALI exposure). (**B**) AECs were treated with IFNs or media control for 6 days during ALI exposure, and mRNA expression of basal cell gene *Krt5* was determined by qPCR (n = 3 for all conditions). (**C**) WT AEC cultures were treated for 6 days with IFN or media control during ALI exposure as shown in (A), and multiciliated cells were measured by immunofluorescence staining of acetylated α-tubulin (the whole transwell membrane scans shown are representative of n = 3 samples for all conditions). All data are representative of at least three independent experiments. Data are shown as mean ± s.e.m. and statistical significance was assessed by two-way ANOVA with Dunnett's post-test (B). ns, not significant (P >0.05); $*P \leq 0.05$.

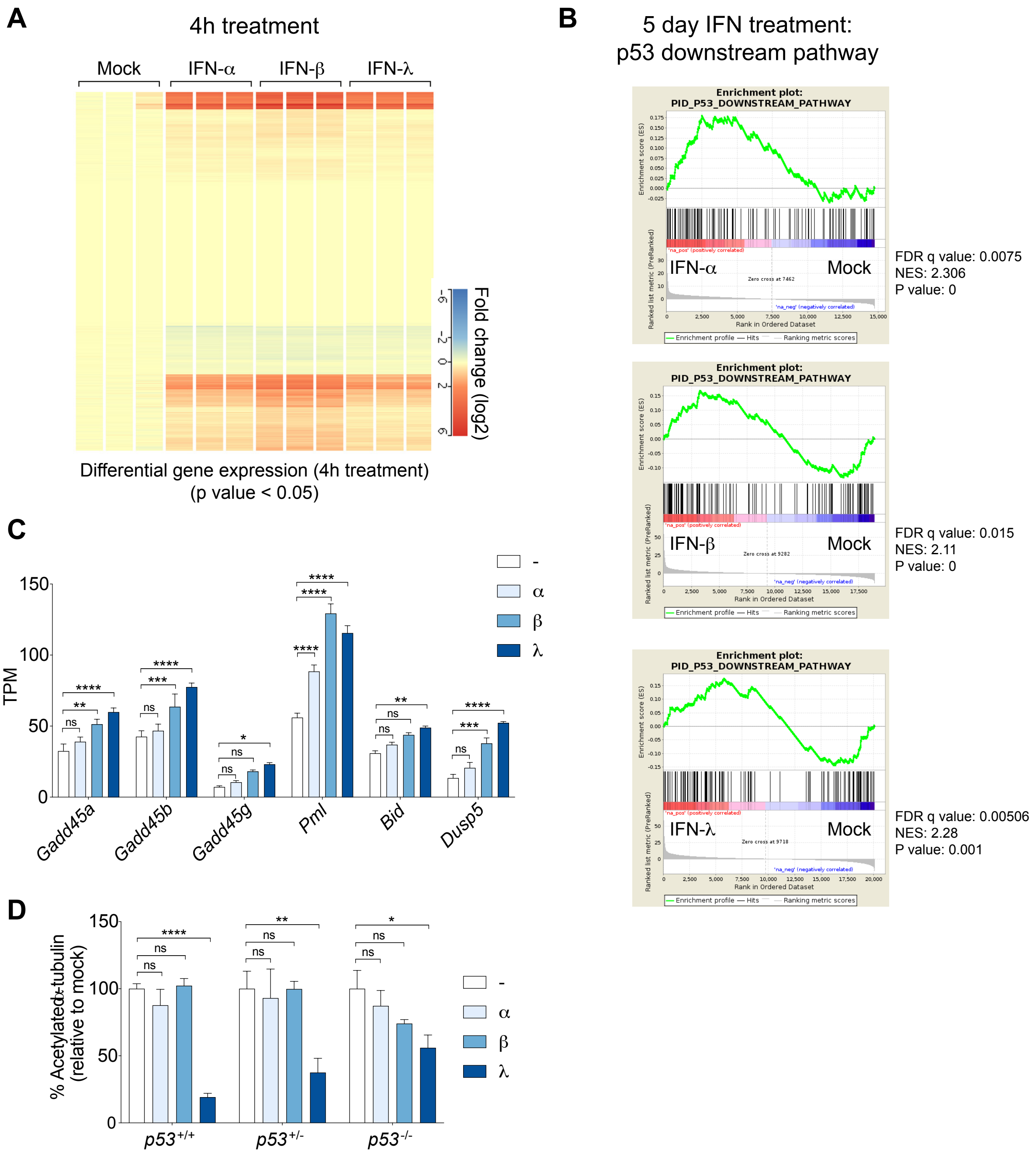


**Fig. S9. p53 pathway and downstream target induction in IFN treated AECs.** (**A**) Heatmap for differentially expressed genes in AECs (one-way ANOVA with Benjamini–Hochberg correction, $P < 0.05$) following 4 hours of IFN treatment, relative to untreated controls (See Data S1 for the full differentially expressed gene list). (**B**) p53 downstream pathway GSEA plots of RNA-seq datasets from 5-day IFN treated cultures, relative to their respective untreated controls. (**C**) Transcripts Per Million (TPM) values for selected p53 downstream target genes. (**D**) Quantification of acetylated α-tubulin fluorescence intensity from entire wells of $p53^{+/+}$, $p53^{+/-}$ and *p53*$^{-/-}$ AEC cultures treated for 6 days with IFN or media control during ALI exposure (n = 3 samples for all conditions). Data are representative of at least two independent experiments (D). Data are shown as mean ± s.e.m. and statistical significance was assessed by two-way ANOVA with Dunnett's post-test. ns, not significant ($P > 0.05$); $*P \leq 0.05$, $**P \leq 0.01$, $***P \leq 0.001$, $****P \leq 0.0001$.

**Table S1: Antibodies and other reagents used for flow cytometry and immunofluorescence**

| Antibody | Clone | Label | Vendor | Dilution |
|---|---|---|---|---|
| EpCam | G8.8 | APC | Invitrogen | 1:800 |
| CD45 | 30-F11 | BV786 | BioLegend | 1:800 |
| CD49f | eBioGoH3 | PE-Cy7 | eBioscience | 1:100 |
| CD31 | RM5228 | BV421 | Invitrogen | 1:800 |
| CD24 | M1/69 | BV510 | BioLegend | 1:200 |
| MHC-II | M5/114.15.2 | BV711 | BioLegend | 1:1000 |
| Ki67 | SolA15 | PE | eBioscience | 1:200 |
| Ly6C | HK1.4 | BV786 | BioLegend | 1:400 |
| Ly6G | 1A8 | FITC | BioLegend | 1:200 |
| CD3 | 17A2 | APC | TONBObiosciences | 1:100 |
| CD19 | 6D5 | APC | BioLegend | 1:100 |
| NK1.1 | PK136 | APC | BioLegend | 1:100 |
| CD64 | X54-5/7.1 | PE-Cy7 | BioLegend | 1:100 |
| Siglec F | E50-2440 | APC-Cy7 | BD Pharmingen | 1:600 |
| CD11b | M1/70 | Pacific blue | BioLegend | 1:200 |
| P53 | GP59-12 | PE | BD Pharmingen | 1:5 |
| Annexin V | N/A | FITC | BioLegend | 1:20 |
| TO-PRO-3 | N/A | APC | Thermofisher | 1:1000 |
| Fixable blue dead stain | BUV395 | UV | Thermo Fisher | 1:400 |
| Acetylated α-tubulin | 6-11B-1 | N/A | Sigma | 1:1000 |

**Data S1. Differential gene expression in 4-hour IFN treated versus mock AECs cultures.** Differentially expressed ($p < 0.05$) genes, 4-hour IFN treatment versus mock AEC cultures (fig. S9A).
